# Corneal lens curvature depends on localized chitin secretion

**DOI:** 10.1101/2025.08.22.671854

**Authors:** Neha Ghosh, Jessica E. Treisman

**Affiliations:** Department of Cell Biology, NYU Grossman School of Medicine, 540 First Avenue, New York, NY 10016

## Abstract

The *Drosophila* corneal lens is an apical extracellular matrix structure with a biconvex shape that enables it to focus light. Here we investigated how this shape is influenced by the source of one of its major components, the polysaccharide chitin. Knocking down the chitin synthase Krotzkopf verkehrt reduced corneal lens thickness and curvature. Conversely, enhancing chitin export by overexpressing Rebuf expanded the corneal lens. We found that the cone and primary pigment cells in the center of each ommatidium produce most of the chitin, and preventing chitin synthesis by these central cells reduced corneal lens curvature. Increasing chitin export from central cells increased the thickness of the central corneal lens, while increasing export from peripheral lattice cells expanded its edges. The biconvex shape thus results from high levels of chitin production by central cells relative to peripheral cells, indicating that localized chitin secretion is critical for normal corneal lens curvature.

## Introduction

The shapes of most biological structures are primarily generated by the numbers, morphologies and positions of their constituent cells. However, extracellular matrix (ECM) can also contribute to tissue and organ morphology. While basement membrane ECM has been shown to provide structural support to many epithelial cell types, the generation of diverse forms from apical ECM (aECM) is less well understood ^1^. aECM structures include chitin-rich arthropod and collagen-rich nematode cuticles which act as permeability barriers and protect against dehydration, luminal scaffolds in tubular organs like the *Drosophila* trachea and *C. elegans* excretory system, mechanosensory structures such as arthropod tactile hairs and mammalian whiskers, and the tectorial membrane of the vertebrate inner ear ^2-5^.

One aECM component that has been shown to play an important role in imparting shape to biological structures is the polysaccharide chitin, a polymer of N-acetylglucosamine ^6^. For instance, newly synthesized chitin is required for the luminal expansion of the fly embryonic tracheal system; in its absence, tracheal tubes form constrictions and cysts instead of expanding uniformly ^7,8^. In *C. elegans*, a chitin layer provides structural rigidity to the eggshell, maintaining the oval shape of the embryo ^9^. Chitin organization can be modified to form structures as diverse as lobster claws and structurally colored butterfly wing scales ^10-12^. Chitin is a major component of insect corneal lenses ^13-15^, but its importance for the morphogenesis of these precisely curved structures is unknown.

The *Drosophila* corneal lens is an aECM structure primarily composed of chitin and chitin-binding proteins arranged in a biconvex shape that is essential to its function of focusing light onto the underlying photoreceptors. We have shown that corneal lens shape is compromised in mutants lacking the Zona Pellucida (ZP) domain-containing proteins Dusky-like (Dyl) or Dumpy (Dpy), and this shape change correlates with a delay in chitin deposition ^16^. In addition, the external corneal lens surface is flattened in mutants lacking the transcription factor Blimp-1, which regulates the expression of genes that encode chitin processing enzymes such as Krotzkopf verkehrt (Kkv), Knickkopf (Knk), Rebuf (Reb), Expansion (Exp), Chitinase 5, Chitinase 7 and Chitinase 10 ^17-21^. These observations prompted us to investigate the role of chitin in creating corneal lens shape.

In this study we show that differential levels of chitin production between the central and peripheral cells of the ommatidium dictate the precise biconvex shape of the *Drosophila* corneal lens. Although all non-neuronal retinal cells express the chitin synthase gene *kkv*, the central cone and primary pigment cells are the major source of chitin for the thick middle region of the corneal lens, while the peripheral lattice cells contribute a small amount of chitin to the tapered corneal lens edges. We showed that increasing chitin deposition by central cells increased the thickness of the middle region, forming spherical corneal lenses. In contrast, increasing chitin deposition by lattice cells made the edges thicker, giving the corneal lenses a rectangular shape. These observations show that localized secretion and restricted diffusion of chitin establish the shape of the corneal lens.

## Results and Discussion

### Chitin accumulates over central cells in the mid-pupal retina

To understand the role of chitin in corneal lens structure, we first characterized its production during corneal lens morphogenesis. Using a fluorescently labeled chitin-binding domain probe (CBD) ^22^ and super-resolution LIGHTNING microscopy, we examined the localization and organization of chitin in mid-pupal retinas. Chitin was not yet present at 48 h after puparium formation (APF) (**Fig. 1A**). Chitin fibers first appeared over primary pigment cells at 50 h APF (**Fig. 1B**), expanding to cone cells and becoming radially arranged at 51 h APF (**Fig. 1C**), and were straighter and more tightly packed at 52 h APF (**Fig. 1D**). By 54 h APF, chitin was evenly distributed in a dense meshwork covering the apical surfaces of the cone and primary pigment cells (**Fig. 1E**). However, no chitin accumulation was observed over the secondary and tertiary pigment cells (lattice cells) at these stages. We examined chitin in horizontal cryosections of late pupal and adult corneal lenses using Calcofluor White staining, which penetrated these dense structures better than the CBD probe ^16,21,23^. At 72 h APF, the corneal lens was externally curved ^16^ and had multiple chitin layers, especially in its central region (**Fig. 1F**). The layers were more numerous and distinct in adult biconvex corneal lenses (**Fig. 1G**). Initial chitin deposition thus occurs only over primary pigment cells and cone cells (collectively called central cells) during mid-pupal stages, and expansion of the proximal-distal axis of the corneal lens involves the progressive addition of chitin layers (**Fig. 1H**).

**Figure 1.**
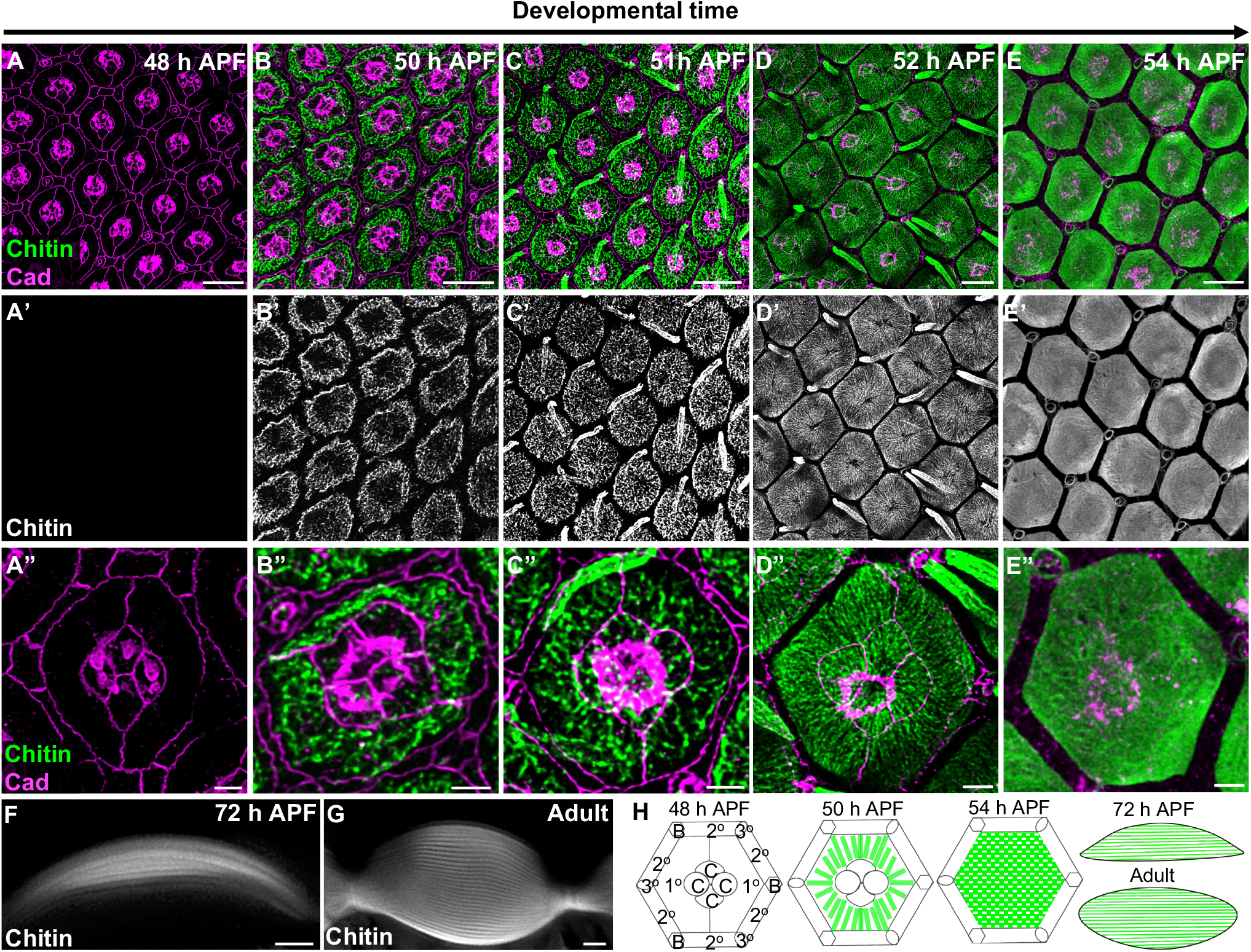
Chitin organization during corneal lens morphogenesis. (**A-E**) Apical surfaces of wild-type (*white*^*1118*^) mid-pupal retinas stained for chitin with CBD (**A’-E’**, green in **A-E**) and E-Cad and N-Cad (Cad, magenta) at the indicated developmental time points. (**A’’-E’’**) show enlargements of single ommatidia. (**F, G**) Horizontal cryosections of wild-type retinas showing 72 h APF (**F**) and adult (**G**) corneal lenses stained for chitin with Calcofluor White. Scale bars: 10 µm (**A-E**), 2 µm (**A”-E”, F-G**). (**H**) Diagram of chitin (green) organization during corneal lens development.

### Disrupting chitin secretion alters corneal lens shape

We next investigated how chitin contributes to corneal lens architecture. The chitin synthase Kkv is an oligomeric transmembrane protein that polymerizes chitin chains and translocates them across the plasma membrane through a central channel ^23,24^ (**Fig. 2A**). We blocked chitin synthesis by generating clones that expressed *kkv* RNAi with *lGMR-GAL4*, which is active in all retinal cells except the mechanosensory bristles ^25^. Kkv antibody staining showed that RNAi expression effectively removed Kkv from the apical surfaces of non-neuronal retinal cells (**Fig. 2B**). Clones expressing *kkv RNAi* showed no apically deposited chitin at 54 h APF, except in the bristles (**Fig. 2C**), indicating that chitin is autonomously synthesized by underlying retinal cells and does not diffuse laterally. Adult corneal lenses produced by *kkv* knockdown ommatidia were completely devoid of chitin and were about half the thickness of wild-type corneal lenses, with reduced external and internal curvature (**Fig. 2D, E, L, M, Fig. 3K**). Interestingly, a significant portion of the corneal lens was still present in the absence of chitin, perhaps because other corneal lens components can be retained by the external scaffold established by the ZP domain-containing proteins Dyl, Dpy and Piopio ^16^. In addition, the outermost layer of the corneal lens is composed of the Retinin protein and waxes ^26^, which may not require chitin for their localization.

**Figure 2.**
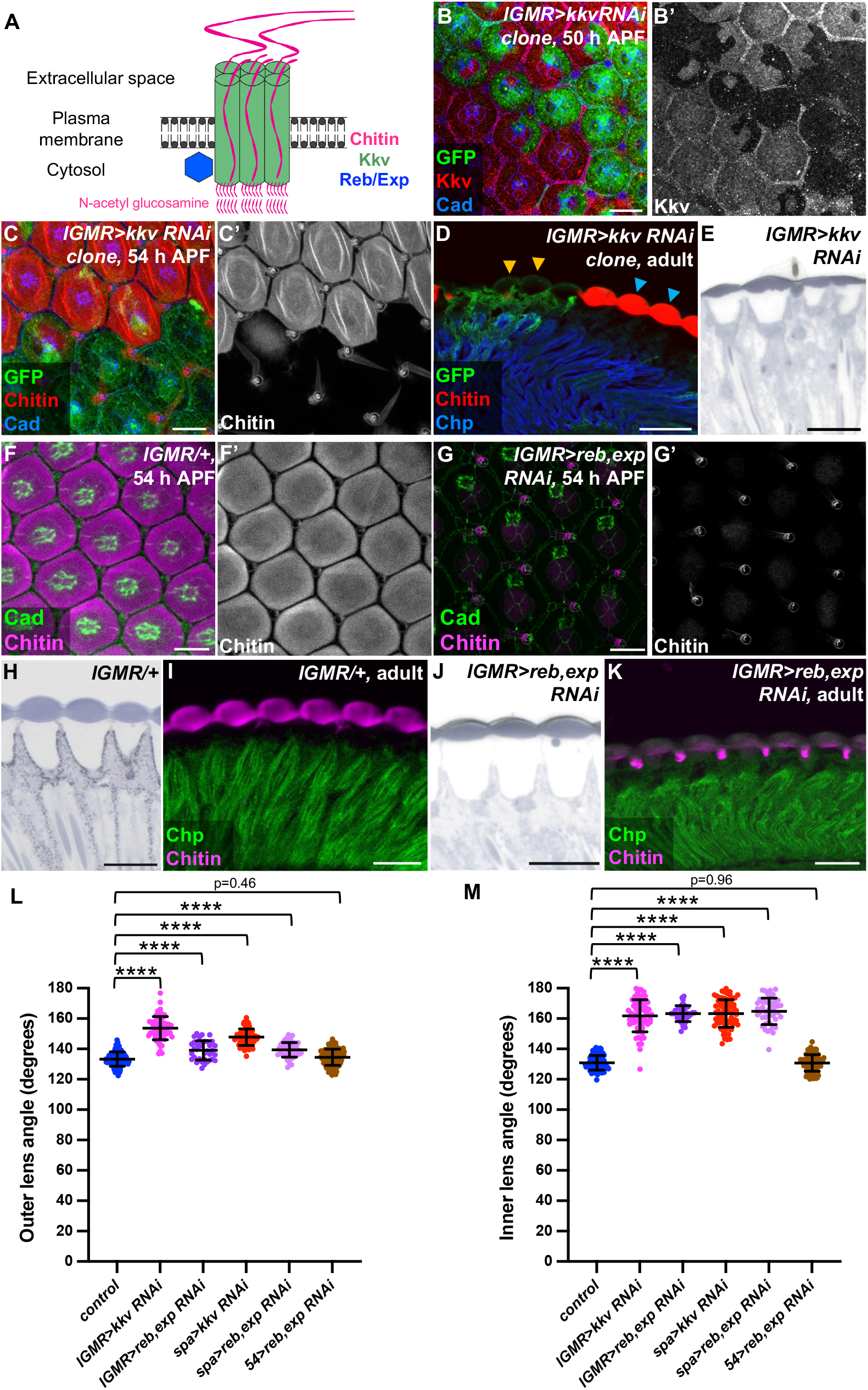
Chitin production is necessary for biconvex corneal lens shape. (**A**) Schematic showing the *Drosophila* chitin production machinery. (**B-D**) Retinas containing clones expressing *kkv* RNAi with *lGMR-GAL4* marked by GFP (green) and stained for Cad (blue, **B, C**) or Chaoptin (Chp, blue, **D**) and Kkv (**B’**, red in **B**) or chitin (**C’**, red in **C, D**). (**B**) At 50 h APF Kkv is lost in *kkv* RNAi-expressing cells. (**C**) at 54 h APF and (**D**) in adults chitin is lost from the corneal lenses. Cyan arrows mark wild-type ommatidia, and yellow arrows mark *kkv* knockdown clones (**D**). (**E**) Plastic section of an *lGMR-GAL4>kkv RNAi* adult eye showing thin corneal lenses. (**F, G**) 54 h APF retinas stained for chitin (**F’, G’**, magenta in **F, G**) and Cad (green). (**F**) control *lGMR-GAL4/+*; (**G**) *lGMR-GAL4*>*reb RNAi; exp RNAi*. (**H-K**) Horizontal sections of control retinas (**H, I**) or *lGMR-GAL4*>*reb RNAi; exp RNAi* retinas (**J, K**). (**H, J**) show plastic sections and (**I, K**) cryosections stained for chitin (magenta) and Chp (green). Scale bars: 10 µm (**B, C, F, G**), 20 µm (**D, E, H-K**). (**L-M**) Graphs showing the outer (**L**) and inner (**M**) angles between adjacent corneal lenses in adult eye sections for wild-type control (n=96 ommatidia/11 retinas), *lGMR>kkv RNAi* (n=64/4), *lGMR>reb RNAi; exp RNAi* (n=35/3), *spa>kkv RNAi* (n=71/4), *spa>reb RNAi; exp RNAi* (n= 47/4), and *54>reb RNAi; exp RNAi* (n=113/9). For all graphs unless otherwise stated, error bars show mean ± SD, and p values were calculated with unpaired two-tailed *t* tests with Welch’s correction and Bonferroni’s correction for multiple comparisons. ^****^, p<0.0001. See also Fig. S1.

**Figure 3.**
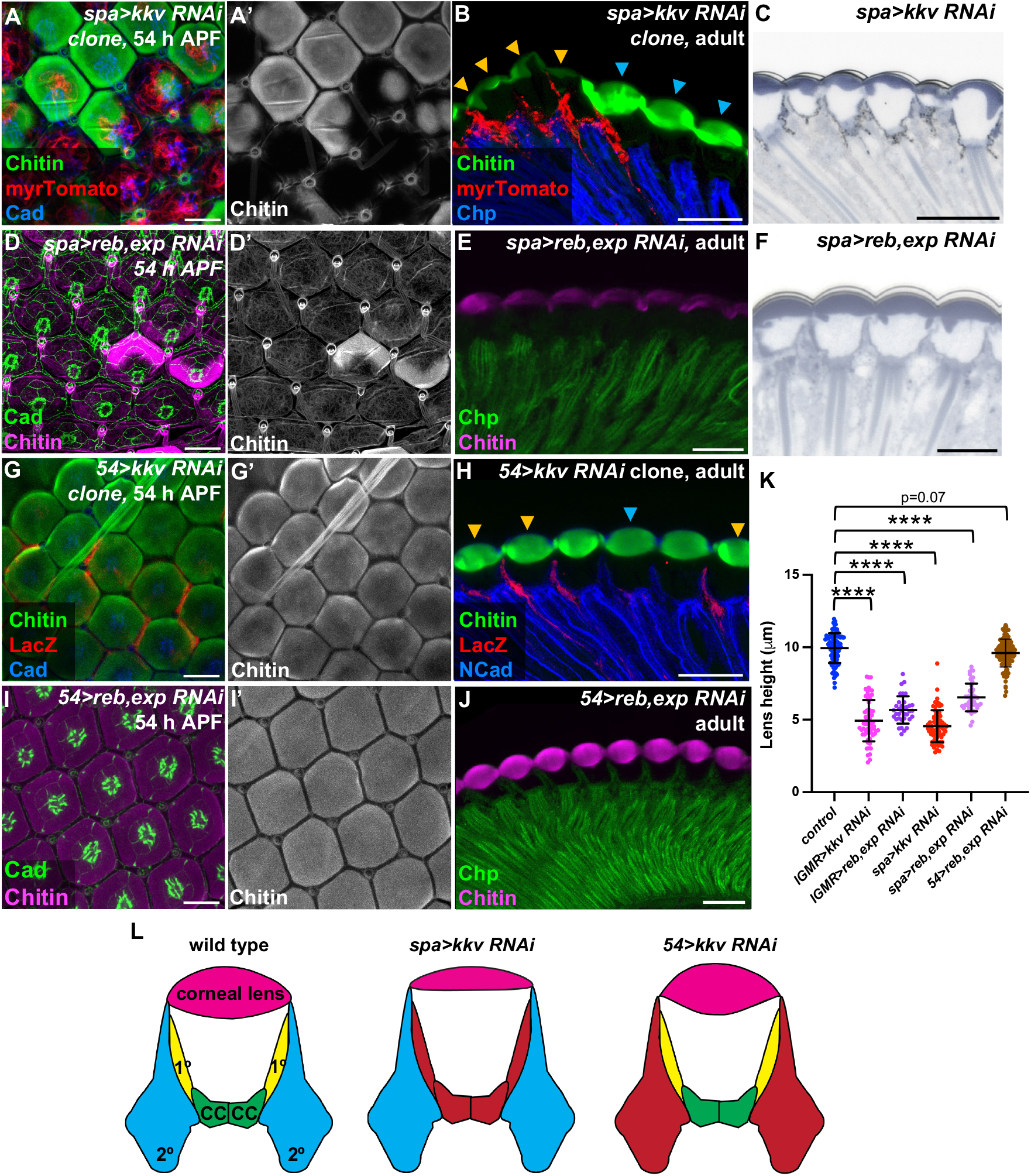
Central cells are the major contributors of chitin. (**A, D, G, I**) 54 h APF retinas; (**B, C, E, F, H, J**) adult eye sections. (**A, B**) *kkv* RNAi expression in central cells with *spa-GAL4* in clones labelled with myrTomato (red), stained for chitin (**A’**, green in **A, B**) and Cad (blue, **A**) or Chp (blue, **B**). (**D, E**) *spa-GAL4>reb* RNAi, *exp* RNAi retinas stained for chitin (**D’**, magenta in **D, E**) and Cad (green, **D**) or Chp (green, **E**). (**C, F**) Plastic sections of *spa-GAL4>kkv RNAi* (**C**) or *spa-GAL4>reb RNAi, exp RNAi* (**F**) adult retinas. (**G-J**) Retinas in which the lattice cell driver *54-GAL4* drives *kkv RNAi* in clones marked with anti-β-galactosidase (red, **G, H**) or *reb* RNAi and *exp* RNAi throughout the retina (**I, J**), stained for chitin (**G’, I’**, green in **G, H**, magenta in **I, J**), and Cad (blue in **G**, green in **I**), NCad (blue, **H**) or Chp (green, **J**). Scale bars: 10 µm (**A, D, G, I**), 20 µm (**B, C, E, F, H, J**). Cyan arrows, wild-type ommatidia; yellow arrows, *kkv* knockdown clones (**B, H**). (**K**) Graph showing the maximum height of corneal lenses in adult eye sections for wild-type control (n=96/11), *lGMR>kkv RNAi* (n=64/4), *lGMR>reb RNAi; exp RNAi* (n=35/3), *spa>kkv RNAi* (n=71/4), *spa>reb RNAi; exp RNAi* (n= 47/4), and *54>rebRNAi; exp RNAi* (n=113/9). (**L**)Model depicting the effects of removing chitin from specific cell types. Red indicates the cells that express *kkv* RNAi. See also Fig. S2.

The chitin translocation activity of Kkv is supported by two interchangeable proteins encoded by the *reb* and *exp* genes ^19,27^ (**Fig. 2A**). Reb and Exp contain Nα-MH2 domains and belong to the atypical SMAD group of proteins ^28-30^. In the retina, Reb expression was observed primarily in lattice cells at 50 h APF (**Fig. S1A**), while Exp was detected only in primary pigment cells at the same stage (**Fig. S1C**), and both could be effectively depleted by RNAi (**Fig. S1B, D**). By 54 h APF, Reb and Exp were both present in all central cells (**Fig. S1E, F**). Knocking down either *reb* or *exp* alone does not affect chitin levels in the embryonic trachea ^19,27^. In the retina, *reb* knockdown with *lGMR-GAL4* did not alter chitin levels at 54 h APF or in adult corneal lenses (**Fig. S1G, H**). Although *lGMR-GAL4*-driven *exp* RNAi caused a loss of apical chitin at 54 h APF, (**Fig. S1I**), the corneal lens showed normal chitin accumulation by the adult stage and its shape was only slightly altered (**Fig. S1J-M**). This suggests that Exp is the primary regulator of chitin secretion in the mid-pupal retina, while Reb compensates for loss of Exp at later developmental stages (**Fig. S1E, F**). Knocking down both *reb* and *exp* in all retinal cells significantly reduced apical chitin deposition in both the 54 h APF retina and the adult corneal lens (**Fig. 2F-K**). Knockdown of *reb* and *exp* resulted in corneal lenses with reduced height and decreased external and internal curvature, effects similar to those caused by depletion of *kkv* (**Fig. 3K, Fig. 2L, M**). Overall, these findings demonstrate that chitin constitutes and/or retains a significant portion of the volume of the corneal lens, consistent with a measurement of 20% chitin content by weight in the dragonfly corneal lens ^13^.

### Chitin production is primarily required in the central cells

As we observed initial chitin deposition only over the central cells, we hypothesized that these cells would be a major source of corneal lens chitin. Indeed, expressing *kkv* RNAi exclusively in clones of central cells using *sparkling-GAL4* (*spa-GAL4*) ^31^ eliminated nearly all chitin over the ommatidia at 50 h and 54 h APF (**Fig. S2A; Fig. 3A**). A small amount of chitin remained in adult corneal lenses, which were abnormally shaped (**Fig. 3B, C, K; Fig. 2L, M**). In contrast, expressing *kkv* RNAi in clones of lattice cells using *54-GAL4* ^32^ did not affect chitin production at 50 h APF (**Fig. S2B**) or its apical accumulation at 54 h APF (**Fig. 3G**). There was a significant reduction in chitin intensity at the edges of the adult corneal lens (**Fig. 3H: Fig. S2D**), suggesting that lattice cells do secrete some chitin later in development. We also tested the effects of blocking chitin translocation by knocking down *reb* and *exp* in either central or lattice cells. Knocking down both *reb* and *exp* in central cells significantly reduced apical chitin deposition at 54 h APF (**Fig. 3D**) and in adult corneal lenses, resulting in thinner corneal lenses with reduced internal and external curvature (**Fig. 2L, M; Fig. 3E, F, K**). However, *reb* and *exp* double knockdown in lattice cells only reduced chitin intensity at the edges of the adult corneal lens, and had no significant impact on earlier chitin accumulation or on corneal lens shape (**Fig. 2L, M; Fig. 3I-K; Fig. S2C-E**). Central cells thus appear to be the primary chitin-producing cells in the retina, and chitin production by these cell types is essential for normal corneal lens size and shape (**Fig. 3L**).

### Localized production of excess chitin alters corneal lens shape

We next investigated the consequences of excessive chitin production. Ectopic *reb* is sufficient to increase chitin deposition by *kkv-*expressing ectodermal cells ^19 27^. We found that overexpressing *reb* in clones of all retinal cells with *lGMR-GAL4* led to premature production of disorganized chitin fibers at 48 h APF (**Fig. 4A**) and a large excess of apical chitin accumulation by 54 h APF (**Fig. 4B**). Adult corneal lenses had distorted shapes, with chitin appearing to leak into the underlying pseudocone (**Fig. 4C**). Chitin entered the pseudocone early in its development, at 84 h APF, and lamellar chitin organization in corneal lenses was disrupted (**Fig. S3A**). To determine how corneal lens shape was affected by the cellular source of excess chitin, we overexpressed *UAS-reb* in clones of either central or lattice cells. *reb* overexpression in central cells caused premature chitin deposition at 48 h APF and excessive chitin accumulation in the center of corneal lenses at 54 h APF (**Fig. 4D, E**). The chitin fibers formed tufts that extended apically, instead of being organized radially. In contrast, overexpressing *reb* in lattice cells resulted in very modest increases in chitin at 48 h and 54 h APF (**Fig. 4G, H**), perhaps because Reb is already present in wild-type lattice cells at 50 h APF (**Fig. S1A**). Adult corneal lens shape was altered by both manipulations. Clones in which central cells misexpressed *reb* produced spherically shaped corneal lenses with increased height, reduced width, and increased outer and inner curvature (**Fig. 4F, J-M**). In contrast, clones in which lattice cells overexpressed *reb* produced rectangular corneal lenses with expanded rather than tapered edges and flattened internal surfaces (**Fig. 4I-M**). Thus, our results suggest that corneal lens shape is determined by differential production of chitin by central and lattice cells. There appears to be little lateral mobility of chitin, so high levels of chitin production by central cells expand the central corneal lens, while lower levels of chitin produced by lattice cells later in development create the tapered corneal lens periphery. Excessive chitin from either source can alter adult corneal lens curvature (**Fig. 4N**).

**Figure 4.**
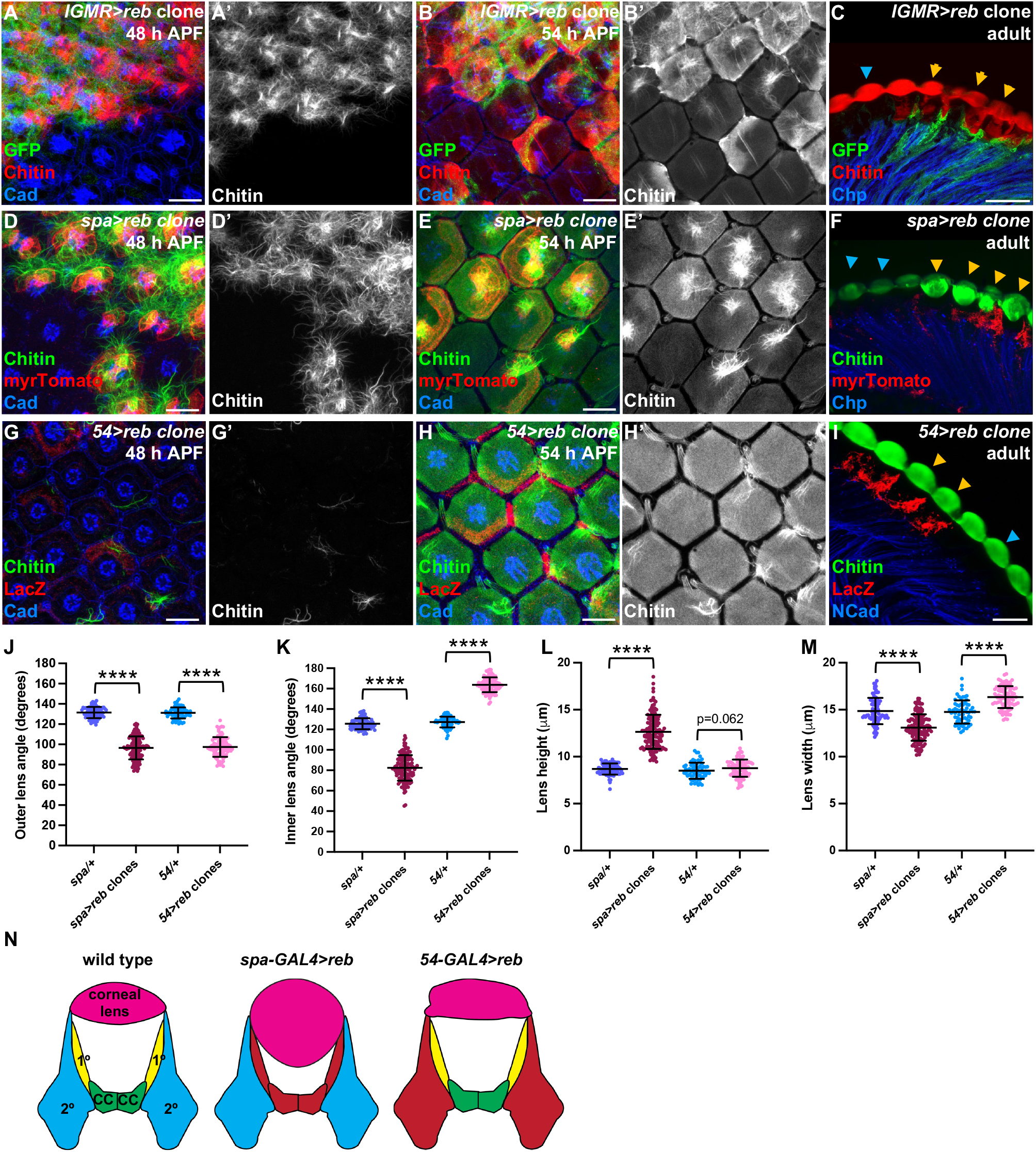
Overproduction of chitin by specific cells alters corneal lens shape. (**A-C**) Misexpression of *reb* with *lGMR-GAL4* in clones marked by GFP (green). (**D-F**) Misexpression of *reb* with *spa-GAL4* in clones marked by myrTomato (red). (**G-I**) Misexpression of *reb* with *54-GAL4* in clones marked by *lacZ* (red). Retinas are stained for chitin (**A’, B’, D’, E’, G’, H’**, red in **A-C**, green in **D-I**) and Cad (blue in **A, B, D, E, G, H**), Chp (blue in **C, F**), or Ncad (blue in **I**). (**A, D, G**) 48 h APF, (**B, E, H**) 54 h APF, (**C, F, I**) adult. Scale bars: 10 µm (**A-B, D-E, G-H**), 20 µm (**C, F, I**). Cyan arrows mark wild-type ommatidia and yellow arrows *reb*-expressing clones (**C, F, I**). (**J-M**) Graphs showing the outer (**J**) and inner (**K**) angles between adjacent corneal lenses, corneal lens height (**L**) and corneal lens width (**M**) in adult eye horizontal cryosections for wild-type control regions and *UAS-reb* clones driven by *spa-GAL4* (n=79/16 wild-type, 122/16 *spa-GAL4>UAS-reb*) or *54-GAL4* (n=63/12 wild-type, 92/12 *54-GAL4>UAS-reb*). (**N**) Model showing the effect of *reb* misexpression in specific retinal cell types on corneal lens structure. Red indicates the cells that express *reb*. See also Fig. S3.

### Controlling chitin secretion and mobility can impart shape to aECM structures

Our study illustrates how differential production of the polysaccharide chitin by specific cell types can sculpt an aECM structure. The cone and primary pigment cells are specialized to secrete high chitin levels necessary for the increased height of the central corneal lens, while chitin production by peripheral lattice cells is more limited and occurs later in development, producing the narrower edges of the corneal lens. Even when produced in vast excess, chitin is unable to spread to neighboring cells, indicating that it must be effectively trapped by plasma membrane and/or ECM proteins. We previously showed that expansion of the apical surfaces of central cells is necessary for them to produce or retain chitin early in pupal development ^16^, suggesting that the narrow apical surfaces of lattice cells might restrict their ability to deposit chitin. Consistent with this possibility, the cells that produce chitinous scales in Manduca undergo endoreplication to expand their size ^33^. Apical expansion of central cells into a domed shape ^34^ may also initiate the curvature of the external corneal lens surface, which is still partially retained even in the absence of chitin. Our observations extend previous work showing that localized secretion contributes to the morphogenesis of other apical ECM structures such as the taenidial folds of the *Drosophila* trachea and the tectorial membrane of the mammalian inner ear ^35-37^.

Chitin fibers can undergo self-assembly into higher-order crystalline structures stabilized by non-covalent interactions ^6,38^. In the insect cuticle, microfibrils consisting of 17-20 antiparallel α-chitin polymers are aligned to form horizontal laminae ^39^. Our observations confirm that the *Drosophila* corneal lens is also constructed of chitin microfibrils that are stacked into distinct layers to expand its height. These layers are less well resolved at the edges of the corneal lens, where they may be thinner and/or more tightly packed. The changes in corneal lens shape that we observe on modulating chitin deposition by cone and primary pigment cells imply that the thickness of the central corneal lens directly depends on the amount of chitin produced by these cells. It is possible that layer thickness or the initiation of new layers is regulated by the local concentration of chitin. Lattice cells are less effective than central cells at contributing chitin to the corneal lens, perhaps due to their lack of *exp* and only transient expression of *reb* at mid-pupal stages. Alternatively, the paucity of chitin deposition by lattice cells might be due to higher expression of chitinases or lower expression of proteins such as Knk that can protect chitin from degradation ^40^. The packing of chitin layers could also be influenced by Knk or other chitin-binding proteins. Mass spectrometric analysis of the fly corneal lens identified only four major proteins ^41^, but similar studies in mosquito ^42^ and transcriptomic analyses of the pupal and adult retina ^21,43^ suggest that the corneal lens contains many additional cuticle proteins which may organize chitin and limit its lateral diffusion.

Due to its nontoxic and biodegradable properties chitin has found numerous applications in biomedicine. For instance, chitin-containing hydrogels are used in corneal transplants ^44,45^ and as dressings to promote wound healing ^46,47^, deacetylated chitin (chitosan) nanoparticles constitute safe drug delivery systems ^48^, and nanocomposite fibers containing chitin are used as scaffolds for bone engineering ^49^. Understanding how chitin scaffolds for biological structures such as the corneal lens are established may provide insights relevant to these biotechnological applications of chitin ^50^.

## Materials and Methods

### Key resources table

| REAGENT or RESOURCE | SOURCE | IDENTIFIER |
| --- | --- | --- |
| **Antibodies** |  |  |
| Chicken polyclonal anti-GFP | Thermo Fisher Scientific | Cat# A-10262<br>RRID: AB_2534023 |
| Rabbit polyclonal anti-DsRed | Takara Bio | Cat# 632496<br>RRID: AB_10013483 |
| Chicken polyclonal anti-β-galactosidase | Abcam | Cat# ab9361<br>RRID: AB_307210 |
| Rat monoclonal anti-ECad | Developmental Studies<br>Hybridoma Bank (DSHB) | #DCAD2<br>RRID: AB_528120 |
| Rat monoclonal anti-NCad | DSHB | #MNCD2<br>RRID: AB_528119 |
| Mouse monoclonal anti-Chp | DSHB | #24B10<br>RRID: AB_528161 |
| Mouse monoclonal anti-Arm | DSHB | #N2 7A1<br>RRID: AB_528089 |
| Rabbit polyclonal anti-Kkv | <sup>27</sup> | N/A |
| Rabbit polyclonal anti-Reb | <sup>19,27</sup> | N/A |
| Rat monoclonal anti-Exp | <sup>19,27</sup> | N/A |
| Goat anti-chicken 488 | Invitrogen | A-11039<br>RRID: AB_2534096 |
| Donkey anti-chicken Cy3 | Jackson ImmunoResearch | 703-165-155<br>RRID: AB_2340363 |
| Goat anti-rabbit Cy3 | Jackson ImmunoResearch | 111-165-003<br>RRID: AB_2338000 |
| Donkey anti-mouse 488 | Invitrogen | A-21202<br>AB_2534069 |
| Donkey anti-mouse Cy3 | Jackson ImmunoResearch | 715-165-151<br>RRID: AB_2315777 |
| Donkey anti-rat Cy3 | Jackson ImmunoResearch | 712-165-150<br>RRID: AB_2340666 |
| Donkey anti-rat Cy5 | Jackson ImmunoResearch | 712-175-153<br>RRID: AB_2340672 |
| **Chemicals, peptides, and recombinant proteins** |  |  |
| Chitin Binding Domain-Alexa Fluor 488 | <sup>16,21</sup> | N/A |
| Chitin Binding Domain-Alexa Fluor 546 | <sup>16,21</sup> | N/A |
| Calcofluor White (Fluorescent Brightener 28 disodium salt solution) | Sigma Aldrich | Cat# 910090 |
| 10% Formaldehyde | Polysciences | Cat# 04018-1 |
| 20% Paraformaldehyde | Fisher Scientific | Cat# 50-980-493 |
| Bovine Serum Albumin (BSA) | Millipore Sigma | A4737 |
| Tissue Freezing Medium (TFM) | GeneralData | TFM-C |
| **Experimental models: Organisms/strains** |  |  |
| *D. melanogaster*: *w*<sup>1118</sup> | Bloomington Drosophila Stock Center (BDSC) | BDSC:3605; FlyBase: FBst0003605 |
| *D. melanogaster*: *IGMR-GAL4* | <sup>25</sup> | N/A |
| *D. melanogaster*: *spa-GAL4* | <sup>31</sup> | BDSC:26656;<br>FlyBase: FBal0127274 |
| *D. melanogaster*: *54-GAL4* | <sup>32</sup> | Flybase: FBti0012688 |
| *D. melanogaster*: *kkv RNAi*<br>*TRiP.HMC05880* | BDSC | BDSC:65006;<br>FlyBase: FBti0184093 |
| *D. melanogaster*: *reb RNAi*<br>*TRiP.HMJ21908* | BDSC | BDSC:57861;<br>FlyBase: FBti0164346 |
| *D. melanogaster*: *exp RNAi*<br>*TRiP.HMS01445* | BDSC | BDSC:35032;<br>FlyBase: FBti0144968 |
| *D. melanogaster*: *UAS-reb* | <sup>27</sup> | Flybase: FBal0302994 |
| *D. melanogaster*: *UAS-CD8-GFP*,<br>*ey3.5-FLP*; *IGMR-GAL4*; *FRT82*, *tub-GAL80/SM6-TM6B* | <sup>21</sup> | N/A |
| *D. melanogaster*: *ey-FLP*, *spa-GAL4*; *UAS-myr-Tomato*; *FRT82*, *tub-GAL80* | <sup>21</sup> | N/A |
| *D. melanogaster*: *UAS-CD8-GFP*, *ey3.5-FLP*; *54-GAL4*, *UAS-lacZ*; *FRT82*, *tub-GAL80/SM6-TM6B* | <sup>21</sup> | N/A |
| *D. melanogaster*: *kkvRNAi*; *FRT82B* | This study | N/A |
| *D. melanogaster*: *rebRNAi*; *exp RNAi* | This study | N/A |
| *D. melanogaster*: *UAS-reb*; *FRT82B* | This study | N/A |
| **Software and algorithms** |  |  |
| Image J | National Institutes of Health | http://imagej.nih.gov.ij/ |
| Adobe Photoshop | Adobe |  |
| GraphPad Prism 10.0 | GraphPad Software | https://www.graphpad.com/ |
| MS-Office (MS-Word, MS-Powerpoint, MS-Excel) | MS-Office |  |
| **Other** |  |  |
| Confocal Microscope | Leica SP8 II |  |
| Super resolution LIGHTNING Microscope | Leica Stellaris 8 |  |

### Fly stocks and genetics

*Drosophila melanogaster* strains were maintained on standard yeast-cornmeal-agar media and raised at 25°C. For analysis of the pupal retina, white prepupae (0 h APF) were collected with a soft wetted brush and cultured at 25°C till the appropriate developmental stage. Both sexes were used interchangeably for all the experiments, as no sex-specific differences were observed.

*Drosophila melanogaster* stocks used to generate *kkv* knockdown clones were: (1) *kkv RNAi TRiP*.*HMC05880; FRT82B/SM6*.*TM6B*; (2) *UAS-CD8-GFP, ey3*.*5-FLP; lGMR-GAL4; FRT82, tub-GAL80/SM6-TM6B* (3) *ey-FLP, spa-GAL4; UAS*-*myr-Tomato*; *FRT82, tub-GAL80* (4) *UAS-CD8-GFP, ey3*.*5-FLP; 54-GAL4, UAS-lacZ; FRT82, tub-GAL80/SM6-TM6B*. Stocks used to knock down *reb* and/or *exp* were: (1) *reb RNAi TRiP*.*HMJ21908*; (2) *exp RNAi TRiP*.*HMS01445*; (3) *reb RNAi; exp RNAi*; (4) *lGMR-GAL4*; (5) *spa-GAL4*; (6) *54-GAL4, UAS-lacZ/ SM6-TM6B*. Stocks used for *reb* overexpression in clones were: (1) *UAS-reb, FRT82B*; (2) *UAS-CD8-GFP, ey3*.*5-FLP; lGMR-GAL4; FRT82, tub-GAL80/SM6-TM6B* (3) *ey-FLP, spa-GAL4; UAS*-*myr-Tomato*; *FRT82, tub-GAL80* (4) *UAS-CD8-GFP, ey3*.*5-FLP; 54-GAL4, UAS-lacZ; FRT82, tub-GAL80/SM6-TM6B*.

Stocks were obtained from the following sources: *UAS-reb* ^27^, *kkv RNAi* (BL65006), *reb RNAi* (BL57861), *exp RNAi* (BL35032): Bloomington *Drosophila* Stock Center (BDSC).

### Immunohistochemistry

For cryosectioning, adult or pupal heads with the proboscis removed were glued onto glass rods using nail polish and fixed for 4 h in 4% formaldehyde in 0.2 M sodium phosphate buffer (pH 7.2) (PB) at 4°C. The heads were then incubated through an increasing sucrose gradient in PB (5%, 10%, 25%, and 30% sucrose) for 20 min each, transferred to plastic molds containing TFM compound and frozen on dry ice. Cryosections of 12 μm were cut at −21°C, transferred onto positively charged slides and postfixed in 0.5% formaldehyde in PB at room temperature (RT) for 30 mins. The slides were then washed in PBS with 0.3% Triton X-100 (PBT) three times for 10 min each, blocked for 1 h at RT in 1% bovine serum albumin (BSA) in PBT and incubated in primary antibodies overnight at 4°C in 1% BSA in PBT. After three 20-min washes in PBT, slides were incubated in secondary antibodies in 1% BSA in PBT for 2 h at RT and mounted in Fluoromount-G (Southern Biotech). A 1:10 dilution of Calcofluor White solution (25% in water; Sigma Aldrich, 910090) was included with the secondary antibodies where indicated.

Pupal retinas attached to the brain were dissected from staged pupae and collected in ice-cold PBS in a glass plate. These samples were fixed on ice in 4% formaldehyde in PBS for 30 min. The samples were washed three times for 10 min each in PBT and incubated overnight at 4°C in primary antibodies in 10% donkey serum in PBT. After three 20-min washes in PBT, the samples were incubated for 2 h in secondary antibodies in PBT/10% serum at RT and washed again three times for 20 min in PBT. Finally, the retinas were separated from the brain and mounted in 80% glycerol in PBS.

The primary antibodies used were: mouse anti-Chp (1:50; Developmental Studies Hybridoma Bank (DSHB), 24B10), chicken anti-GFP (1:400; Thermo Fisher, A-10262), chicken anti-LacZ (1:1000; Abcam, ab9361), rat anti-Ecad (1:10, DSHB, DCAD2), rat anti-Ncad (1:50, DSHB, DN-Ex), rabbit anti-Kkv (1:300)^27^, rabbit anti-Reb (1:100)^19,27^, rat anti-Exp (1:100) ^19,27^. All antibodies were validated either using mutant or knockdown conditions as shown or by verifying that the staining pattern matched previously published descriptions. The secondary antibodies used were from either Jackson ImmunoResearch (Cy3 or Cy5 conjugates used at 1:200) or Invitrogen (Alexa488 conjugates used at 1:1000). Fluorescently labeled SNAP-CBD-probes (1:200) ^16,21^ were included with the secondary antibodies. Images were acquired on a Leica SP8 confocal microscope for normal confocal microscopy or on a Leica Stellaris for super-resolution LIGHTNING microscopy with a 63X oil immersion lens and processed using ImageJ and Adobe Photoshop.

### Plastic sections

Adult heads of *Drosophila* were dissected and fixed in a freshly made fixative containing 2% paraformaldehyde, 2.5% glutaraldehyde and 0.05% Triton X-100 in 0.1 M sodium cacodylate buffer (pH 7.2) at room temperature for 4 h and then overnight at 4^º^C. The fixed heads were rinsed with 0.1 M sodium cacodylate buffer and post-fixed with 1% OsO_4_ in 0.1 M cacodylate buffer, followed by dehydration in a graded ethanol series (30%, 50%, 70%, 85%, 95%, 100%), infiltrated with propylene oxide/Spurr mixtures and finally embedded in Spurr resin (Electron Microscopy Sciences, PA, USA). Semithin horizontal 1 mm sections were cut and mounted on a glass slide, then baked on a hot plate overnight at 37 ^º^ C. The sections were stained with 0.1% Toluidine Blue, dried on a hot plate and imaged with a Zeiss Axioplan microscope and processed in Adobe Photoshop.

### Quantification and statistical analysis

The outer and inner angles between adjacent corneal lenses were measured according to the schematic in Fig. S1K, using the angle tool in ImageJ. To measure lens height, freehand straight lines were drawn in the center of the corneal lens from the upper to lower surface using the line tool in ImageJ and measured. Total chitin fluorescence was measured in a rectangle 3 μm wide centered on the junction between two corneal lenses and normalized to the fluorescence in a rectangle of the same width centered on the midpoint of the corneal lens. Values were plotted in GraphPad Prism v10. Significance was calculated using Welch’s two-tailed unpaired t-tests with Bonferroni correction for multiple comparisons when necessary. Sample numbers and definitions of error bars are given in the figure legends.

## Acknowledgments

We thank Marta Llimargas, the Bloomington *Drosophil*a stock center, the Vienna *Drosophila* resource center, the Kyoto stock center, and the Developmental Studies Hybridoma Bank for fly stocks and reagents. Information available on FlyBase was invaluable for this work. We thank NYULH DART Microscopy Laboratory Alice Liang, Jason Yin and Jason Liang for consultation and assistance with plastic sections, and this core is partially funded by NYU Cancer Center Support Grant NIH/NCI P30CA016087. The manuscript was improved by the critical comments of Gira Bhabha, Maria Bustillo, Holger Knaut, Sudershana Nair, and Pragati Sharma. This work was funded by the National Institutes of Health (grant R01 EY035624 to J.E.T.).

## Author contributions

Conceptualization, N.G. and J.E.T.; investigation, N.G.; data curation, N.G.; formal analysis, N.G. and J.E.T.; writing-original draft, N.G.; writing – review and editing, J.E.T.; funding acquisition, J.E.T.; supervision, J.E.T.

