## Supplementary Figures and Legends for "Corneal lens curvature depends on localized chitin secretion"

### Supplementary Information

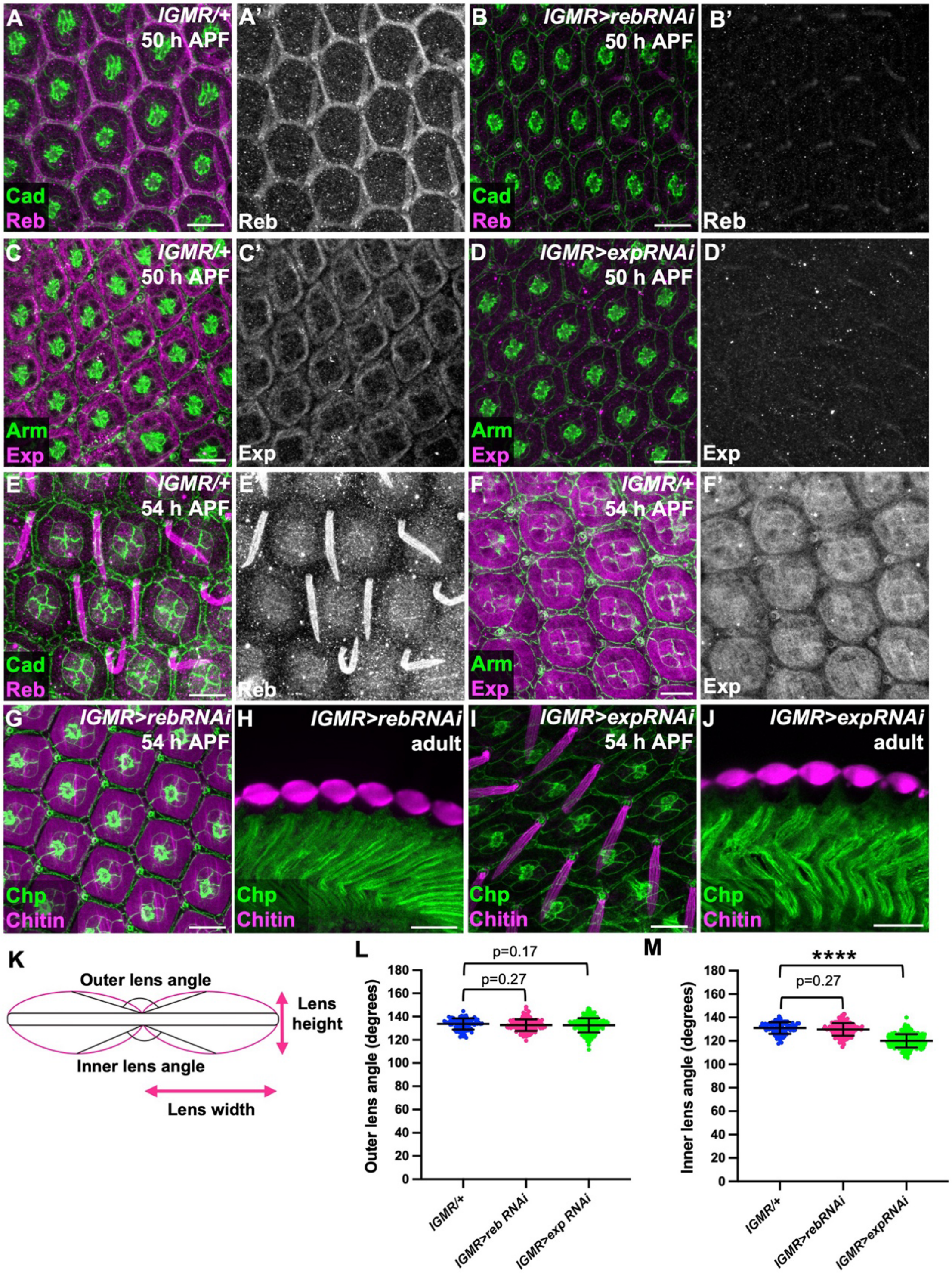


**Figure S1**

**Figure S1. Reb and Exp act redundantly on corneal lens morphology, related to Figure 2.** (**A-F**) Retinas stained for Cad (green) and Reb (**A', B', E'**, magenta in **A, B, E**) or Exp (**C', D', F'**, magenta in **C**, **D, F**). (**A, C, E, F**) control; (**B**) *lGMR>reb RNAi*; (**D**) *lGMR>expRNAi*. (**A-D**), 50 h APF; (**E, F**) 54 h APF.

(**G-J**) *lGMR>rebRNAi* (**G, H**) or *lGMR>expRNAi* (**I, J**) 54 h APF retinas (**G, I**) or horizontal adult eye cryosections (**H, J**) stained for chitin (magenta) and Cad (green in **G, I**) or Chp (green in **H, J**).

Scale bars: 10 μm (**A-F, G, I**), 20 μm (**H, J**).

(**K**) Schematic defining the outer and inner angles between adjacent corneal lenses. The corneal lens height and width are also depicted.

(**L-M**) Graphs showing the outer (**L**) and inner (**M**) angles between adjacent corneal lenses in adult eye horizontal cryosections for wild-type control (n=80/6), *lGMR>reb RNAi* (n=188/13), *lGMR>exp RNAi* (n=134/18).

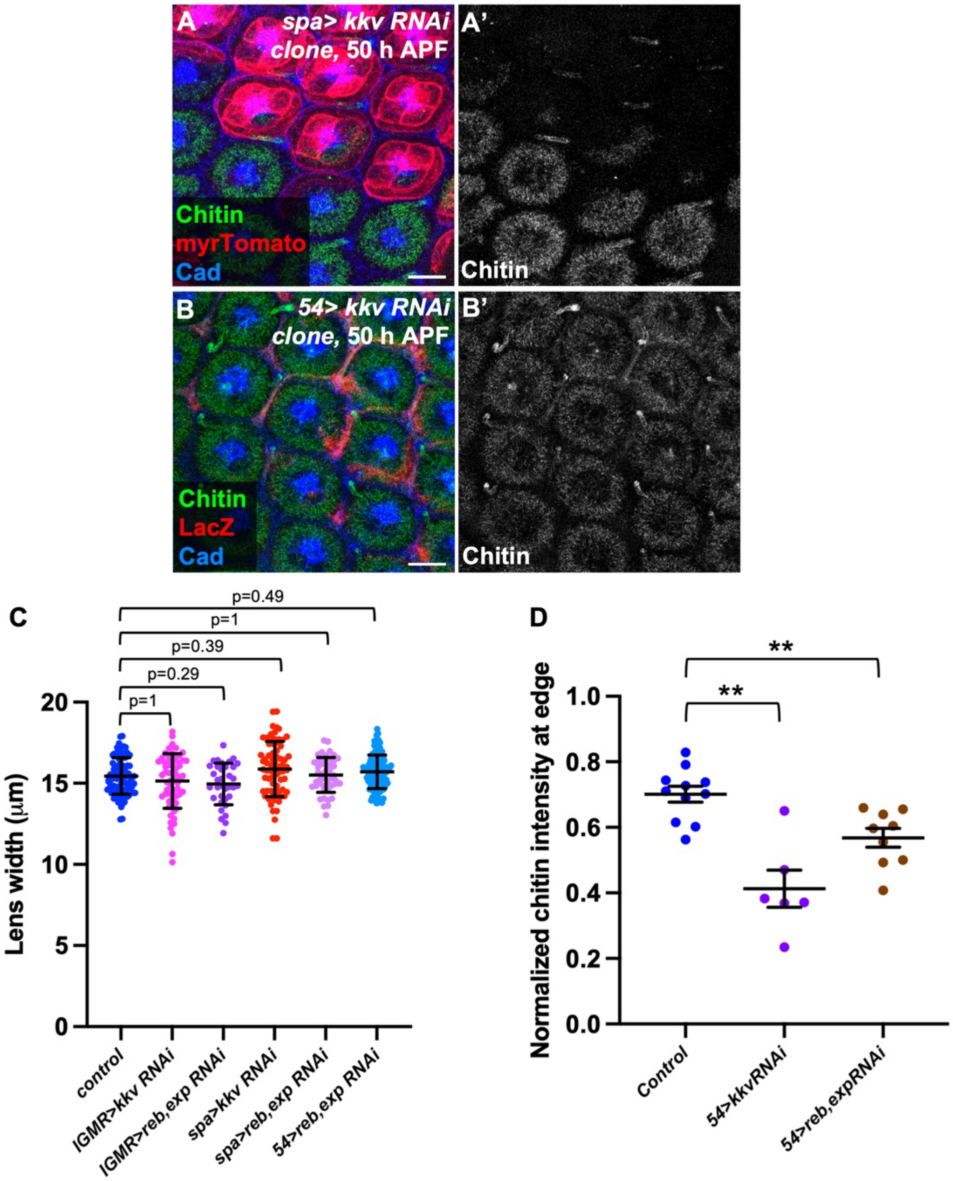


**Figure S2**

**Figure S2. Loss of chitin from lattice cells has a minor effect on corneal lens architecture, related to Figure 3.**

(**A, B**) 50 h APF retinas in which *kkv RNAi* is expressed in central cells with *spa-GAL4* in clones labelled with myrTomato (red, **A**) or in lattice cells with *54-GAL4* in clones labelled with anti-β-galactosidase (red, **B**), stained for chitin (**A'**, **B'**, green) and Cad (blue).

Scale bars: 10 µm.

(**C**) Graph showing corneal lens width in adult plastic sections or cryosections for wild-type control (n=96/11), *IGMR>kkvRNAi* (n=64/4), *IGMR>rebRNAi; expRNAi* (n=35/3)*, spa>kkv RNAi* (n=71/4), *spa>rebRNAi; expRNAi* (n= 47/4) and *54>rebRNAi; exp RNAi* (n=113/9).

(**D**) Graph showing chitin fluorescence intensity in a 3 μm wide region at the adult corneal lens edge normalized to its center in wild-type control (n=79/11), *54>kkv RNAi* clones (n=14/6) and *54>rebRNAi; expRNAi* (n=113/9). **p=0.0048 (*54>kkv RNAi* v. control), **p=0.0052 (*54>reb, exp RNAi* v. control). Each point represents the mean value for one retina, and error bars show mean ± SEM.

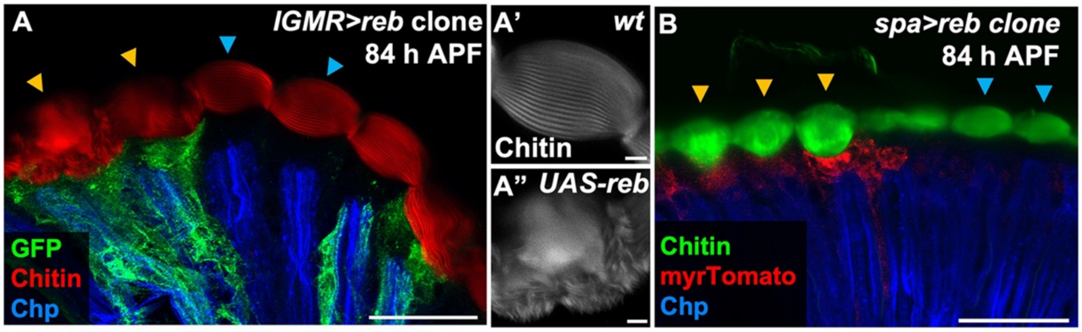


**Figure S3**

**Figure S3. Effect of excess chitin in the late pupal corneal lens, related to Figure 4.**
(**A, B**) Cryosections of 84 h APF retinas containing clones labelled with GFP (green, **A**) or myrTomato (red, **B**) in which *UAS-reb* is driven by *IGMR-GAL4* (**A**) or *spa-GAL4* (**B**), stained for chitin (red in **A**, green in **B**) and Chp (blue). (**A', A"**) show enlargements of individual wild-type (**A')** or *reb*-misexpressing (**A"**) corneal lenses stained for chitin with Calcofluor White. Scale bars, 20 μm (**A, B**), 2 μm (**A', A"**). Cyan arrows mark wild-type ommatidia and yellow arrows *reb*-expressing clones.
